# Conjunctive Targeting Links Drug Synergy to Emergent Proteome Structural States

**DOI:** 10.64898/2026.09.24.752349

**Authors:** Guoqing Cheng, Rabah Soliymani, Elham Gholizadeh, Pouya Behrouzi, Mahdi Muhaddesi, Risto Renkonen, Markku Varjosalo, Amir A. Saei, Mohieddin Jafari

## Abstract

Combinatorial therapies are widely used in the treatment of acute myeloid leukemia (AML) to address disease heterogeneity, adaptive resistance, and rewired signaling and metabolic states. Yet drug prioritization remains largely guided by clinical or phenotypic evidence, while the molecular mechanisms underlying effective drug combinations remain incompletely defined. To narrow this gap, we developed Combinatorial high-ratio Partial proteolysis with reference PRoteome Analysis (CoPPRA), a structural proteomics workflow based on limited proteolysis of cell lysates that profiles drug-associated changes in regional protein accessibility at peptide-level resolution. Here, we applied CoPPRA to ruxolitinib and ulixertinib, individually and in combination, in AML-related cell lysates. Our findings extend conjunctive targeting (CT), a recently proposed mechanism of combinatorial drug action in which combined exposure produces protein targeting patterns not observed with either drug alone. Previously identified through combination-associated changes in protein solubility/stability, CT is examined here at peptide-level resolution through regional differences in proteolytic accessibility. The ruxolitinib–ulixertinib combination produced broad peptide-level accessibility changes, including a subset meeting the predefined criteria for CT. CT candidates predominantly exhibited regional accessibility changes, with altered peptide regions occurring against comparatively small changes across the remaining quantified peptides from the same proteins. MAP2K1 and ATP6V1G1 showed pronounced differences between overlapping peptide sequences, highlighting localized variation in combination-associated accessibility, including an ATP6V1G1 peptide mapping to an annotated helical region. Combination-associated increases in peptide signals were also observed in PIK3R1, BRD4, and PTPN11, linking regional accessibility changes to signaling and transcriptional regulators relevant to AML. Functional enrichment and network analyses further implicated nucleotide and glucose metabolism, ficolin-1-rich granules, ribosome-associated processes, and phagocytic vesicles. These results extend conjunctive targeting from protein-level solubility/stability changes to regional differences in proteolytic accessibility, showing that combination-associated effects can be concentrated within specific peptide regions rather than distributed uniformly across proteins. More broadly, CoPPRA provides a peptide-resolved approach for investigating the molecular features of combinatorial drug action and prioritizing protein regions for subsequent mechanistic validation.

## Introduction

Cancer therapy fails and cancer relapses not only because tumors acquire resistance, but because malignant cells can adapt across multiple biological layers under sustained treatment pressure. In acute myeloid leukemia (AML), cells adapt to therapy through genetic evolution, reversible changes in cell state, and metabolic remodeling^1–3^. Interactions with the bone marrow microenvironment further support leukemic cell survival during treatment^4^. In general, these overlapping survival mechanisms sustain residual disease and contribute to relapse, motivating combination therapies that address multiple vulnerabilities within the leukemic cell population and improve the depth and durability of treatment responses^5,6^.

At the molecular level, however, the widespread use of drug combinations does not by itself explain how combinations overcome adaptive resistance. Ultimately, the mechanisms mentioned above converge on changes in target engagement, signaling propagation, protein conformation, and pathway flux. These changes determine whether a drug can exert durable control over malignant cells. Therefore, drug action is not only a matter of nominal target inhibition but also of how ligand binding reshapes the stability, accessibility, and functional state of proteins within a dynamic cellular network. In resistant systems, however, these effects are often incomplete, redistributed, or rapidly rewired, which makes it difficult to infer the mechanism from the phenotypic response alone. In this case, a mechanistic understanding of combinatorial therapy requires readouts that connect drug exposure to defined molecular interactions and state changes within the proteome^7–9^.

Combination effects are commonly framed in terms of additivity or synergy, that is, whether the joint response exceeds predictions derived from independent action or dose-additivity models^10^. Although useful for quantifying phenotypic interaction, these frameworks do not resolve its molecular basis, thereby limiting mechanistic interpretation and constraining the rational design of drug combinations. Here, we distinguish cooperativity from conjunctive targeting as related but distinct mechanistic concepts. In cooperativity, each agent produces an observable molecular or cellular effect individually, and the combination amplifies or prolongs that effect. In conjunctive targeting, by contrast, the defining feature is the emergence of target proteins or conformational states that are detectable only under the combination and cannot be inferred from either single-agent condition^11^. Although phenotypic synergy is well documented, direct molecular evidence for this combination-specific layer remains limited^6,11,12^, highlighting the need for approaches that can identify conjunctive targets and define their causal sites.

Proteome-wide thermodynamic profiling provides a scalable discovery layer for conjunctive targeting^8,11^. Our lab’s previous workflow, Combinatorial Proteome Integral Solubility/Stability Alteration analysis (CoPISA), quantifies proteome-wide solubility/stability shifts and identifies signatures that emerge specifically under combination drug exposure, providing initial evidence of conjunctive targeting. This concept follows an AND-gate logic, in which a molecular effect emerges when both drugs are present but is not observed with either drug alone^11,13^.^11^. However, these global signatures offer limited insight into the specific protein regions, domains, or structural features underlying combination-associated effects^7,9,11^. Although resolving the precise active conformational states of proteins remains challenging at the proteome-wide scale, identifying changes at the level of protein domains, structural motifs, and functional compartments can provide valuable mechanistic insights into these effects.

To bridge this gap, we developed CoPPRA (Combinatorial high-ratio Partial proteolysis with reference PRoteome Analysis), a peptide-resolved structural proteomics workflow specifically designed to investigate protein accessibility changes associated with drug combinations. CoPPRA builds on established limited proteolysis approaches, including LiP-MS^14^, PELSA^15^, and HOLSER^16^, while adapting their methodological principles to combinatorial drug profiling. Whereas PELSA combines high-ratio partial proteolysis with label-free DIA quantification, HOLSER incorporates fully digested reference samples into a TMT-based multiplexed workflow. CoPPRA combines high-ratio partial proteolysis using a 1:1 trypsin-to-protein ratio with label-free DIA quantification and separately acquired fully digested reference samples. Importantly, the workflow integrates individual and combined drug treatments within a comparative experimental design, enabling the identification of protein regions exhibiting altered proteolytic accessibility under combination exposure. By extending peptide-level structural profiling to combinatorial drug treatments, CoPPRA provides a means of investigating combination-associated proteome remodeling and identifying potential molecular mechanisms underlying drug synergy.

## Results

### CoPPRA delivers deep and reproducible peptide-resolved accessibility profiling in AML lysates

CoPPRA is a structural proteomics workflow with the core idea of partial proteolysis under non-denaturing conditions followed by label-free LC-MS/MS quantification to profile protein accessibility at peptide resolution. Partial proteolysis translates ligand- and state-dependent differences in proteolytic accessibility into quantitative peptide signatures, which enables the detection of treatment-associated structural changes^7,9^. The experimental design also includes fully digested reference samples acquired in separate LC-MS/MS runs, providing a complementary peptide profile to the partially digested treatment samples. Here, we applied CoPPRA to NOMO-1 cell lysates treated with vehicle (DMSO), ruxolitinib, ulixertinib, or their combination. Previous AML combination-therapy studies identified this drug pair as a promising regimen with antileukemic activity and low toxicity^6^ and implicated conjunctive targeting in its effects^11^. These findings motivated its use as a model for investigating combination-dependent changes in protein accessibility.

Briefly, NOMO-1 native lysates were prepared under non-denaturing buffer conditions, exposed to vehicle, ruxolitinib, ulixertinib, or the combination treatment, and subjected to a short pulse of trypsin proteolysis to generate structure-sensitive peptide patterns, followed by label-free LC-MS/MS analysis on Orbitrap platforms (**Fig. 1**). The study included three biological replicates per treatment condition and three fully digested reference samples, yielding 15 samples analyzed across 45 raw files. CoPPRA quantified 4,220 leading protein groups and 42,876 unique peptide sequences, with consistent identification coverage across technical runs and substantial coverage across biological replicates and conditions (**Supplementary Fig. 1A**). The raw-file peptide matrix had an overall missingness of 27.4% for treatment groups and 32.3% in reference groups, while the biological sample peptide matrix had an overall missingness of 12.2% and 17.2% for the treatment and reference groups. This value reflects, in part, differences in peptide detection between the fully digested reference samples and partially digested samples, which generate distinct peptide profiles. To distinguish these expected differences from variability in peptide detection, missingness was additionally evaluated within treatment conditions and across replicates. Proteins supported by at least two distinct peptide sequences were retained, and peptides detected in at least 20% of raw files were kept. Following median normalization, peptide intensity distributions were comparable across samples (**Supplementary Fig. 1B**). The correlation analysis further illustrated quantitative consistency across both technical runs and biological replicates (**Supplementary Fig. 1C**). Collectively, these results demonstrated broad proteomic coverage and satisfactory reproducibility, establishing a reliable dataset for downstream analysis of treatment-associated changes in protein accessibility.

**Fig. 1.**
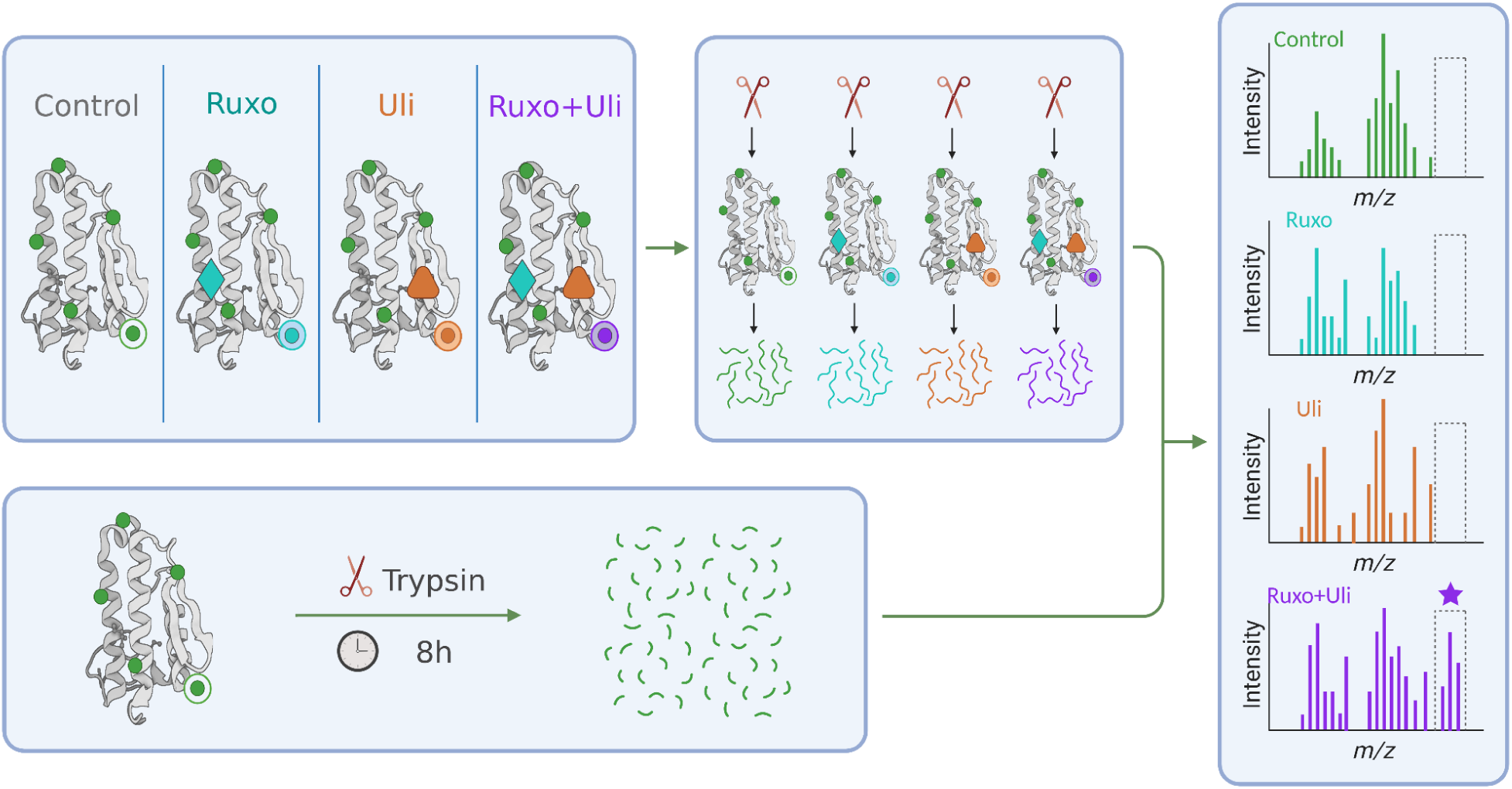
CoPPRA workflow for peptide-resolved profiling of drug-induced structural accessibility. Native NOMO-1 lysates were treated with vehicle control, ruxolitinib (R; labeled Ruxo in the schematic), ulixertinib (U; labeled Uli in the schematic), or their combination (RU) under non-denaturing conditions. Drug- and combination-dependent changes in protein conformation alter local protease accessibility, which is captured by short trypsin proteolysis and converted into treatment-specific peptide patterns. In parallel, fully digested reference samples were analyzed in separate LC-MS/MS runs to support peptide identification. Samples were analyzed individually by label-free DIA LC-MS/MS, and treatment-dependent peptide accessibility changes were quantified to identify proteins and protein regions specifically affected by the drug combination.

### Combination treatment defines a distinct structural accessibility landscape beyond either single agent

Unsupervised analysis of normalized CoPPRA structural-peptide intensities revealed treatment-associated variation in the accessibility landscape. PCA restricted to the 12 vehicle- and drug-treated samples showed that PC1, PC2, and PC3 explained 15.13%, 12.73%, and 11.70% of the variance, respectively. The treatment groups occupied distinct but partially overlapping regions of the PCA space, with their centroids displaced from the vehicle and from one another, particularly in the PC1–PC3 projection (**Fig. 2A**). These patterns indicate treatment-associated differences in peptide-level accessibility, while the within-group dispersion highlights variability among biological replicates. Differential accessibility analysis further supported a combination-enriched structural response. Using a nominal threshold of P < 0.05 and an absolute fold-change cutoff of >2-fold, 2,392 peptides were differentially accessible following ruxolitinib–ulixertinib treatment, compared with 2,518 following ulixertinib and 1,887 following ruxolitinib treatment (**Fig. 2C**). Across the three comparisons, 5,078 distinct peptides met the selection criteria. To assess whether the observed number of significant peptides exceeded that expected under random treatment assignment, sample labels were randomly permuted 10,000 times, retaining three biological samples per condition. The same statistical testing pipeline and selection criteria were applied to each permuted dataset. The observed total number of peptides significant in at least one treatment comparison exceeded the empirical null expectation (global permutation *P* = 0.0013), supporting an overall treatment-associated accessibility response.

**Fig. 2.**
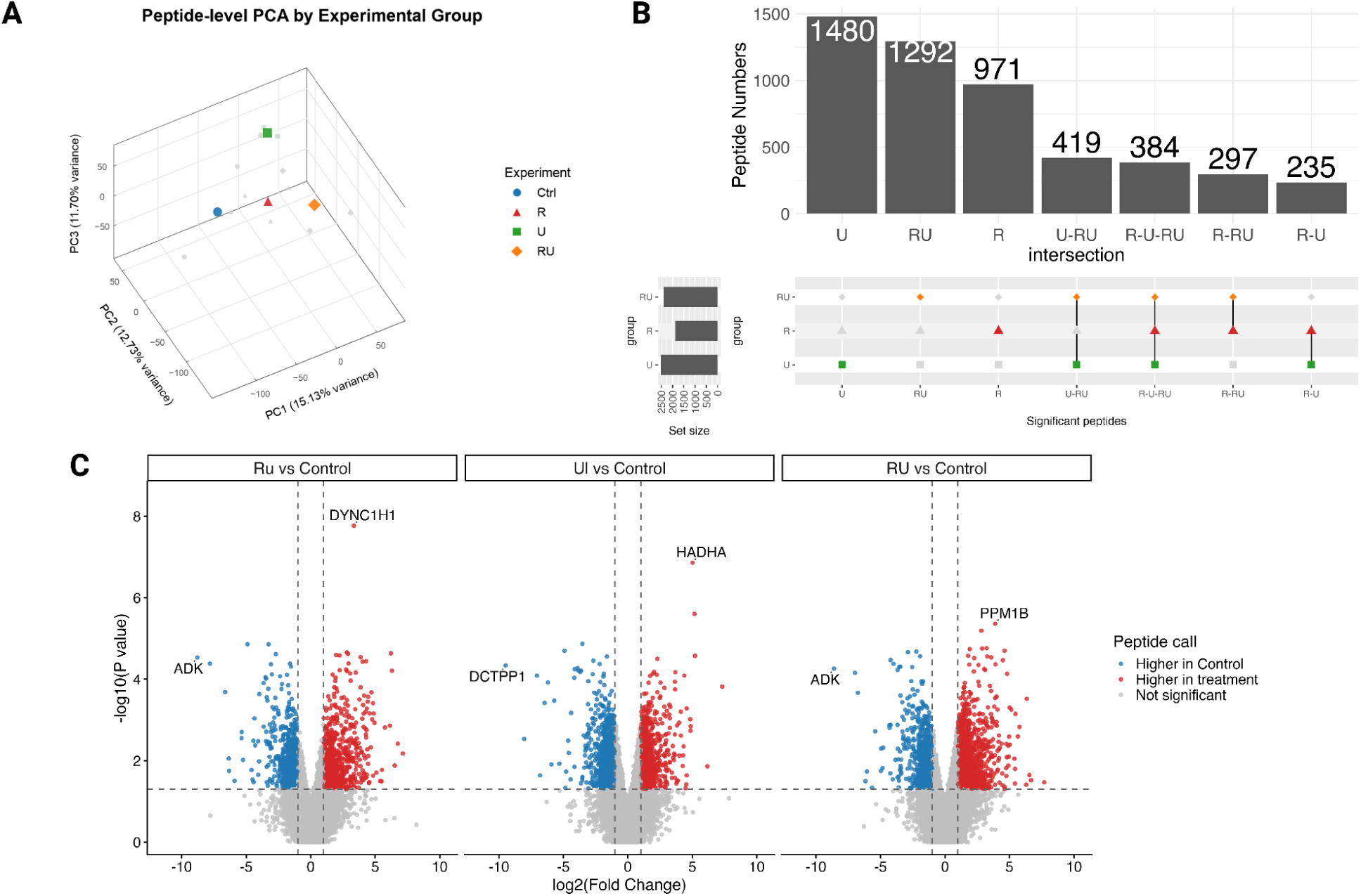
Combination treatment induces a distinct CoPPRA accessibility profile. **A**, Peptide-level PCA of normalized CoPPRA intensities across control, ruxolitinib (R), ulixertinib (U), ruxolitinib plus ulixertinib (RU) and fully digested reference samples. **B,** UpSet analysis of significant peptide-associated accessibility changes across R, U and RU conditions, identifying the RU-specific conjunctive targeting protein set. **C,** Volcano plots showing peptide-level differential accessibility for R, U and RU relative to control. Red and blue points indicate peptides increased in treatment or control, respectively; grey points indicate non-significant peptides. Dashed lines mark the significance and fold-change thresholds (P<0.05 and abs(Log2fc)>1). The permutation analysis for 10,000 iterations generated a global p-value of 0.0013.

Intersection analysis showed that 1,292 RU-responsive peptides met the significance criteria exclusively under combination treatment, representing 54.0% of the RU-responsive peptide set and defining the peptide-level conjunctive targeting set (**Fig. 2B**). A further 419 peptides were shared exclusively between ulixertinib and RU, 297 between ruxolitinib and RU, and 384 across all three treatments. These results identify a substantial combination-specific component of the accessibility response at the applied thresholds. At the protein level, the corresponding analysis identified 306 proteins meeting the conjunctive targeting criterion.

Within the 1,292 conjunctive targeting peptides, five biologically interesting candidates with large accessibility changes and low nominal P values mapped to ASCC2, C1QBP, PIK3R1, BRD4 and MAP2K1. The ASCC2 peptide PALPLDQLQITHKDPK showed a pronounced increase under combination treatment (log2FC = 3.58, *P* = 2.92 × 10⁻⁵), while the PIK3R1 peptide ISEIIDSR increased by log2FC = 3.04 (*P* = 5.65 × 10⁻⁵). The C1QBP peptide EVSFQSTGESEWK exhibited a larger shift (log2FC = 4.00, *P* = 0.00113). Similarly, the BRD4 peptide NSNPDEIEIDFETLKPSTLR and MAP2K1 peptide IPEQILGK increased by log2FC = 2.97 and 4.15, respectively (*P* = 0.00113 and 0.00343). Each peptide met the differential-accessibility criteria under RU treatment, while neither monotherapy comparison met both criteria. These peptide-resolved responses highlight specific regions contributing to the conjunctive targeting profile and prioritize candidates for subsequent structural and functional investigation.

Exact UniProt accession matching between the CoPPRA conjunctive targeting set and the CoPISA NOMO-1 RU AND-gate sets identified 15 shared proteins in intact cells and seven in lysates (Table 1). STOML2 appeared in both comparisons, yielding 21 unique proteins across 22 context-specific matches. All seven lysate matches (OPA1, POLD3, THOC2, SH3BGRL3, PHRF1, UBQLN2 and STOML2) had positive CoPPRA RU-responsive peptide log₂ fold changes (1.00–1.89). For every matched accession, at least one CoPPRA peptide and the corresponding CoPISA protein met the nominal *P* < 0.05 criterion. CoPISA protein-level fold changes also varied in direction within each context. OPA1 and STOML2 are involved in mitochondrial organization, POLD3 in DNA replication and repair, and UBQLN2 in protein quality control, making these proteins biologically distinct representatives of the shared set. CoPPRA and CoPISA values reflect peptide-level and protein-level measurements, respectively.

**Table 1.** Proteins shared between the CoPPRA and CoPISA NOMO-1 lysate conjunctive targeting sets. Exact UniProt accession matching identified 22 proteins shared with the CoPISA RU AND-gate set. CoPPRA values correspond to the listed RU-responsive peptides; CoPISA values are protein-level measurements.

| CoPISA sample | Gene | UniProt | Protein |
| --- | --- | --- | --- |
| Intact cells | ARPC2 | O15144 | Actin-related protein 2/3 complex subunit 2 |
| Intact cells | SERPINB1 | P30740 | Leukocyte elastase inhibitor |
| Intact cells | RPL27A | P46776 | Large ribosomal subunit protein uL15 |
| Intact cells | RPL5 | P46777 | Large ribosomal subunit protein uL18 |
| Intact cells | RPL23A | P62750 | Large ribosomal subunit protein uL23 |
| Intact cells | RPS24 | P62847 | Small ribosomal subunit protein eS24 |
| Intact cells | RPL10A | P62906 | Large ribosomal subunit protein uL1 |
| Intact cells | RNASEH2B | Q5TBB1 | Ribonuclease H2 subunit B |
| Intact cells | SUPV3L1 | Q8IYB8 | Mitochondrial RNA helicase SUPV3L1 |
| Intact cells | GADD45GIP1 | Q8TAE8 | Large ribosomal subunit protein mL64 |
| Intact cells | MRPL44 | Q9H9J2 | Large ribosomal subunit protein mL44 |
| Intact cells | STOML2 | Q9UJZ1 | Mitochondrial stomatin-like protein 2 |
| Intact cells | PADI2 | Q9Y2J8 | Protein-arginine deiminase type 2 |
| Intact cells | TMA7 | Q9Y2S6 | Translation machinery-associated protein 7 |
| Intact cells | SNX9 | Q9Y5X1 | Sorting nexin 9 |
| Lysate | OPA1 | O60313 | Dynamin-like 120 kDa protein |
| Lysate | POLD3 | Q15054 | DNA polymerase delta subunit 3 |
| Lysate | THOC2 | Q8NI27 | THO complex subunit 2 |
| Lysate | SH3BGRL3 | Q9H299 | SH3 domain-binding glutamic acid-rich-like protein 3 |
| Lysate | PHRF1 | Q9P1Y6 | PHD and RING finger domain-containing protein 1 |
| Lysate | UBQLN2 | Q9UHD9 | Ubiquilin 2 |
| Lysate | STOML2 | Q9UJZ1 | Mitochondrial stomatin-like protein 2 |

### Structural-peptide responses recover drug-related signalling proteins and AML-associated candidates

To assess the biological relevance of the accessibility screen, we examined significant peptides from established drug targets and closely connected signalling proteins. The ERK1/MAPK3 peptide LKELIFQETAR decreased under RU treatment (log2FC = −1.75, *P* = 0.0387). The ERK2/MAPK1 peptide GQVFDVGPR increased under ulixertinib (log2FC = 4.11, *P* = 0.000628). Peptides from the upstream kinases MEK1/MAP2K1 (KLEELELDEQQR) and MEK2/MAP2K2 (KLEELELDEQQKK) decreased under ulixertinib (log2FC = −1.36 and −1.18; *P*= 0.0479 and 0.0184, respectively). JAK1/2 was not represented in the tested peptide table, but the JAK–STAT pathway component STAT5A yielded a combination-responsive peptide, AEHQVGEDGFLLK (log2FC = 3.18, *P* = 0.0135). These findings support recovery of proteins associated with the signalling pathways targeted by ulixertinib and ruxolitinib^17,18^.

Within the conjunctive targeting set, several peptides combined large effects with low nominal *P* values. ASCC2 peptide PALPLDQLQITHKDPK and PIK3R1 peptide ISEIIDSR increased under RU treatment (log2FC = 3.58 and 3.04; *P* = 2.92 × 10⁻⁵ and 5.65 × 10⁻⁵, respectively). C1QBP peptide EVSFQSTGESEWK and MAP2K1 peptide IPEQILGK also showed pronounced increases (log2FC = 4.00 and 4.15; *P* = 0.00113 and 0.00343). Each met the combined significance and effect-size criteria under RU, while neither monotherapy comparison met both criteria, identifying peptide-level responses characteristic of conjunctive targeting.

Conjunctive targeting candidates connected the accessibility response to established AML biology. One piece of the most clear evidence is the BRD4 peptide NSNPDEIEIDFETLKPSTLR that met the conjunctional-targeting criterion (log2FC = 2.97, *P* = 0.00113); BRD4 has an experimentally established role in sustaining the proliferation and self-renewal of leukemic cells^19^, identifying an accessibility response in nucleophosmin, a protein recurrently mutated in AML^20^. Additionally, PTPN11/SHP2 peptide GVDGSFLARPSK met the conjunctional-targeting criterion (log2FC = 1.16, *P* = 0.00988), linking the RU response to a RAS signalling regulator recurrently activated by mutations in AML^20,21^.

### MEK1 and ATP6V1G1 show distinct accessibility responses across peptide regions

Peptide-resolved profiles extended the candidate-level analysis by revealing how treatment responses differed within individual proteins. Across nine quantified MEK1/MAP2K1 peptides, RU produced a heterogeneous accessibility profile (**Fig. 3A**). IPEQILGK showed the largest increase under RU (log₂FC = 4.15, *P* = 0.00343), compared with R (0.44, *P* = 0.958) and U (2.33, *P* = 0.100; **Fig. 3B**). Only RU satisfied both the statistical and effect-size thresholds, placing this peptide in the conjunctive targeting set. The overlapping AGRIPEQILGK peptide changed little under RU (log₂FC = 0.19), showing that closely overlapping sequences could yield distinct response profiles. In contrast, KLEELELDEQQR decreased under U (log₂FC = −1.36, *P* = 0.0479), whereas its RU response did not reach nominal significance (−1.11, *P* = 0.113; **Fig. 3C**). The remaining MEK1 peptides also failed to meet both response criteria under RU. This within-protein heterogeneity supports interpreting the MEK1 response at peptide resolution.

**Fig. 3.**
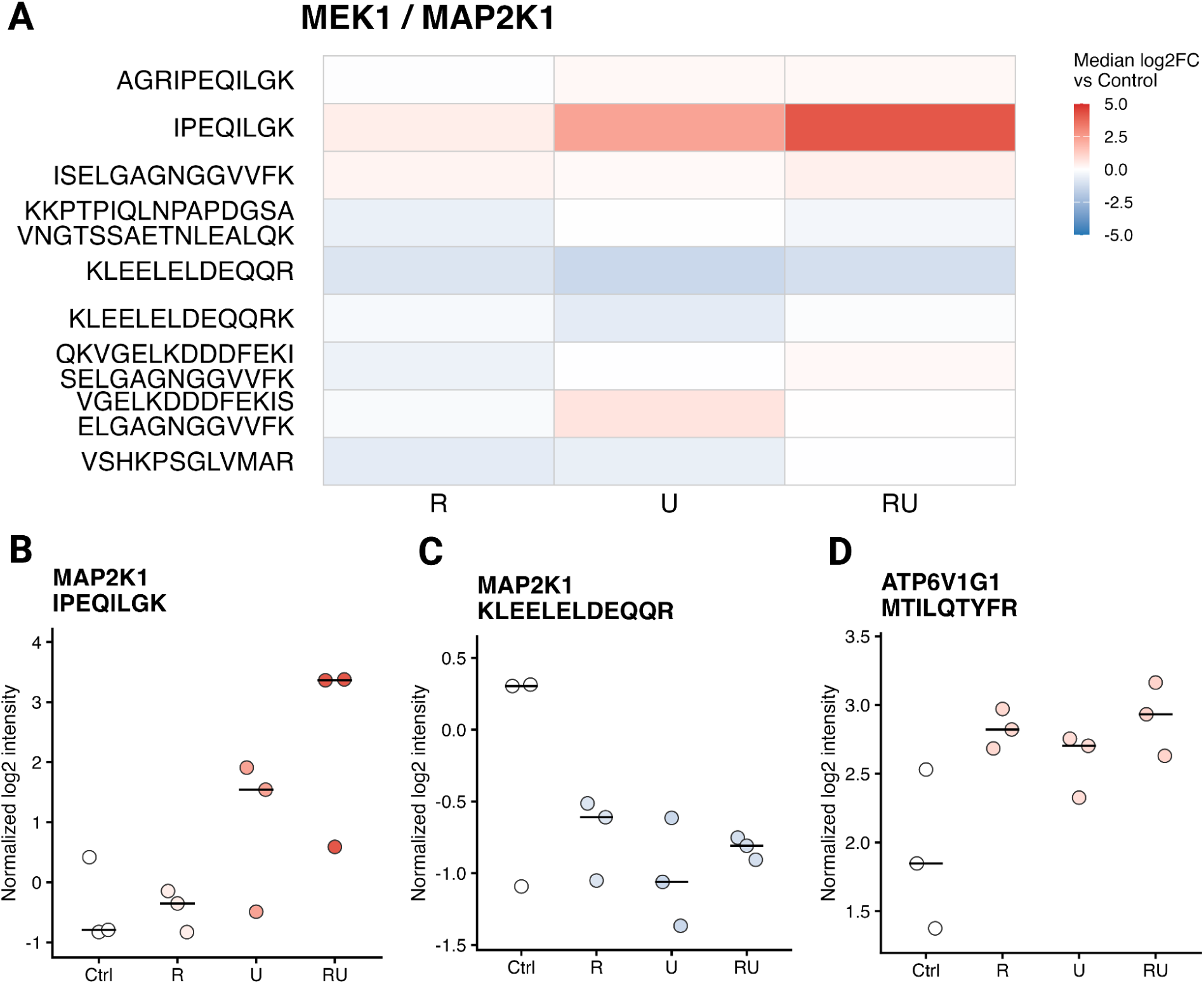
Peptide-resolved accessibility responses of MEK1 and ATP6V1G1. **A,** Heatmap of median log₂ fold changes relative to vehicle control for nine quantified MEK1/MAP2K1 peptides following ruxolitinib (R), ulixertinib (U), or combined treatment (RU). **B–D,** Normalized log₂ intensities for MAP2K1 peptides IPEQILGK (**B**) and KLEELELDEQQR (**C**), and the ATP6V1G1 peptide MTILQTYFR (**D**). Each point represents one biological replicate (n = 3 per condition), with horizontal bars indicating group medians. Colors indicate log₂ fold changes relative to control on a common scale from −5 to 5. Differential accessibility was defined by nominal limma P < 0.05 and |log₂FC| > 1. Ctrl, vehicle control.

The V-ATPase subunit ATP6V1G1 provided a complementary example with sequence-level localization. Its MTILQTYFR peptide increased under RU (log₂FC = 1.09, *P* = 0.0100), with smaller increases under R (0.97, *P* = 0.0161) and U (0.86, *P* = 0.0585; **Fig. 3D**). All three peptides displayed in the replicate plots were measured in three biological replicates per condition. The RU response therefore occurred against a background of detectable, same-direction monotherapy changes.

Mapping all six quantified ATP6V1G1-associated sequences placed MTILQTYFR at residues 81–89, within a UniProt-annotated helical segment (**Fig. 4**). Overlapping peptides spanning residues 33–48 did not meet both response thresholds under any treatment. The N-terminal peptide ASQSQGIQQLLQAEKR also responded under RU (log₂FC = 1.50, *P* = 0.0437), whereas its shorter overlapping sequence, ASQSQGIQQLLQAEK, changed little (log₂FC = 0.06). Both N-terminal sequences also matched ATP6V1G2; MTILQTYFR uniquely matched ATP6V1G1 in the sequence search. Altogether, these findings show how CoPPRA distinguishes localized peptide responses within candidate proteins and provide a sequence-resolved foundation for examining whether these candidates converge on shared biological functions.

**Fig. 4.**
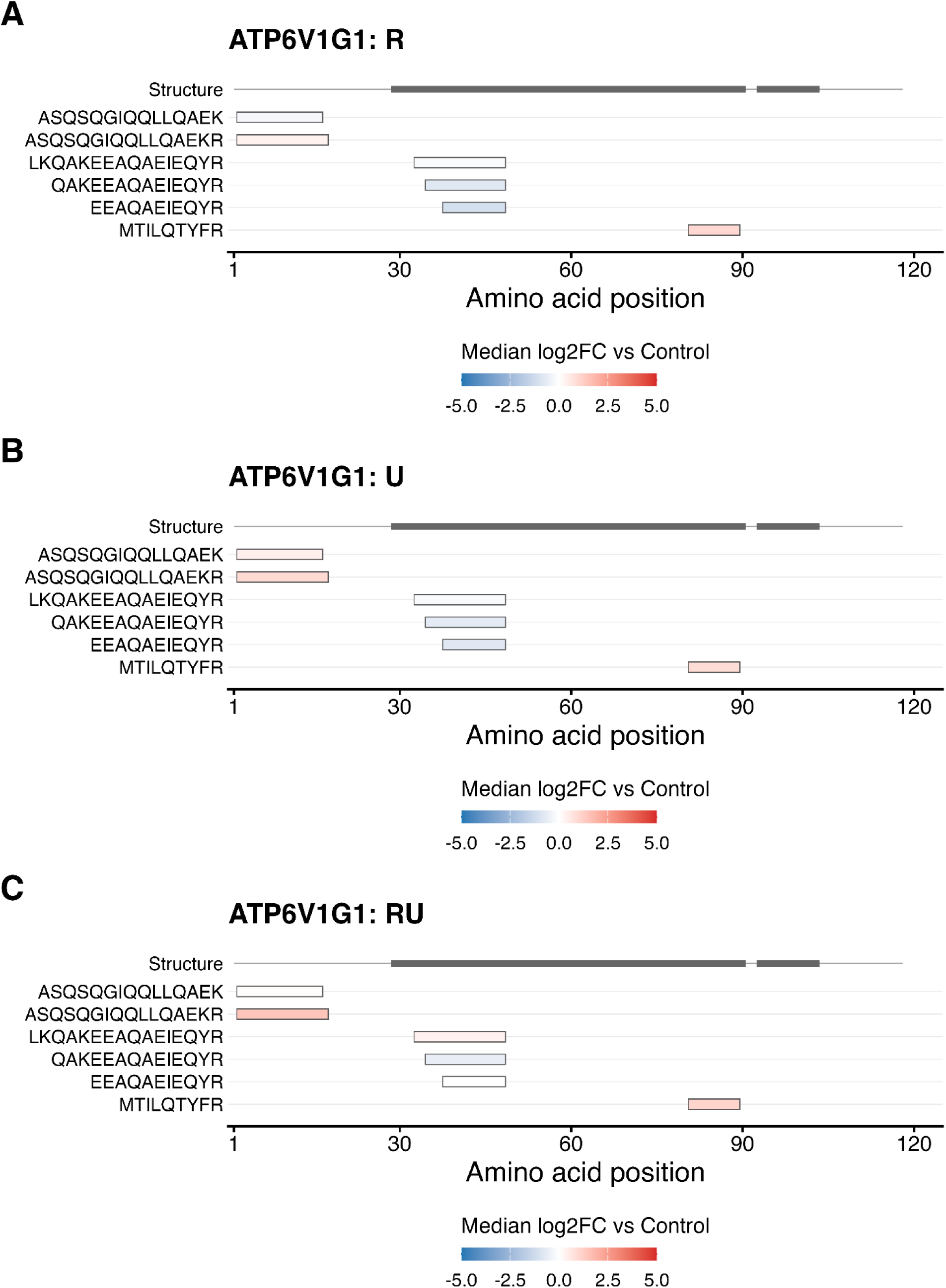
Sequence mapping of ATP6V1G1 peptide accessibility responses. Peptide responses following ruxolitinib (**A**), ulixertinib (**B**), and combined treatment (**C**) mapped onto the ATP6V1G1 sequence (UniProt O75348); colors represent log₂ fold changes relative to vehicle control on a common scale from −5 to 5. Grey bars in the structure track indicate UniProt-annotated α-helices. The MTILQTYFR peptide spans residues 81–89. The two N-terminal sequences, ASQSQGIQQLLQAEK and ASQSQGIQQLLQAEKR, also match ATP6V1G2 and therefore do not uniquely identify ATP6V1G1.

### Enrichment analysis links conjunctive targeting to nucleotide metabolism, ribosome biogenesis, and granule-associated compartments

To determine whether CoPPRA-defined responses converged on shared biological processes, we performed GO and KEGG enrichment analysis across proteins showing accessibility changes following ruxolitinib, ulixertinib, combined treatment, and conjunctive targeting. The single and combined treatment profiles shared strong negative chromatin- and nucleosome-associated signatures. Ruxolitinib additionally showed negative enrichment of megakaryocyte- and myeloid-differentiation-related processes. Ulixertinib retained the chromatin response and extended it to cytoplasmic translation, ribosomal, adhesion-, and junction-associated terms. The full combination profile retained these chromatin- and translation-associated responses. It also added positive enrichment of purine and nucleoside-phosphate metabolism, glycolysis/gluconeogenesis, and nucleotide metabolism, together with negative ribosome-biogenesis terms. Thus, the R, U, and RU profiles indicate a shared chromatin response and broader biosynthetic and metabolic remodeling following combined JAK–ERK inhibition.

Within this broader response, the conjunctive targeting set showed a coordinated shift towards nucleotide metabolism and away from ribosome biogenesis. Biological process analysis linked positive enrichment to nucleotide biosynthesis and metabolism, whereas ribonucleoprotein complex formation, small ribosomal subunit biogenesis, and organelle assembly were negatively enriched (**Fig. 5A**). Cellular component analysis revealed positive enrichment of ficolin-1-rich granule compartments and negative enrichment of ribosome-associated and phagocytic vesicle compartments (**Fig. 5B**). Molecular function analysis further highlighted methyltransferase activity (adjusted *P* = 0.092; **Fig. 5C**), providing additional exploratory evidence of functional alterations associated with treatment.

**Fig. 5.**
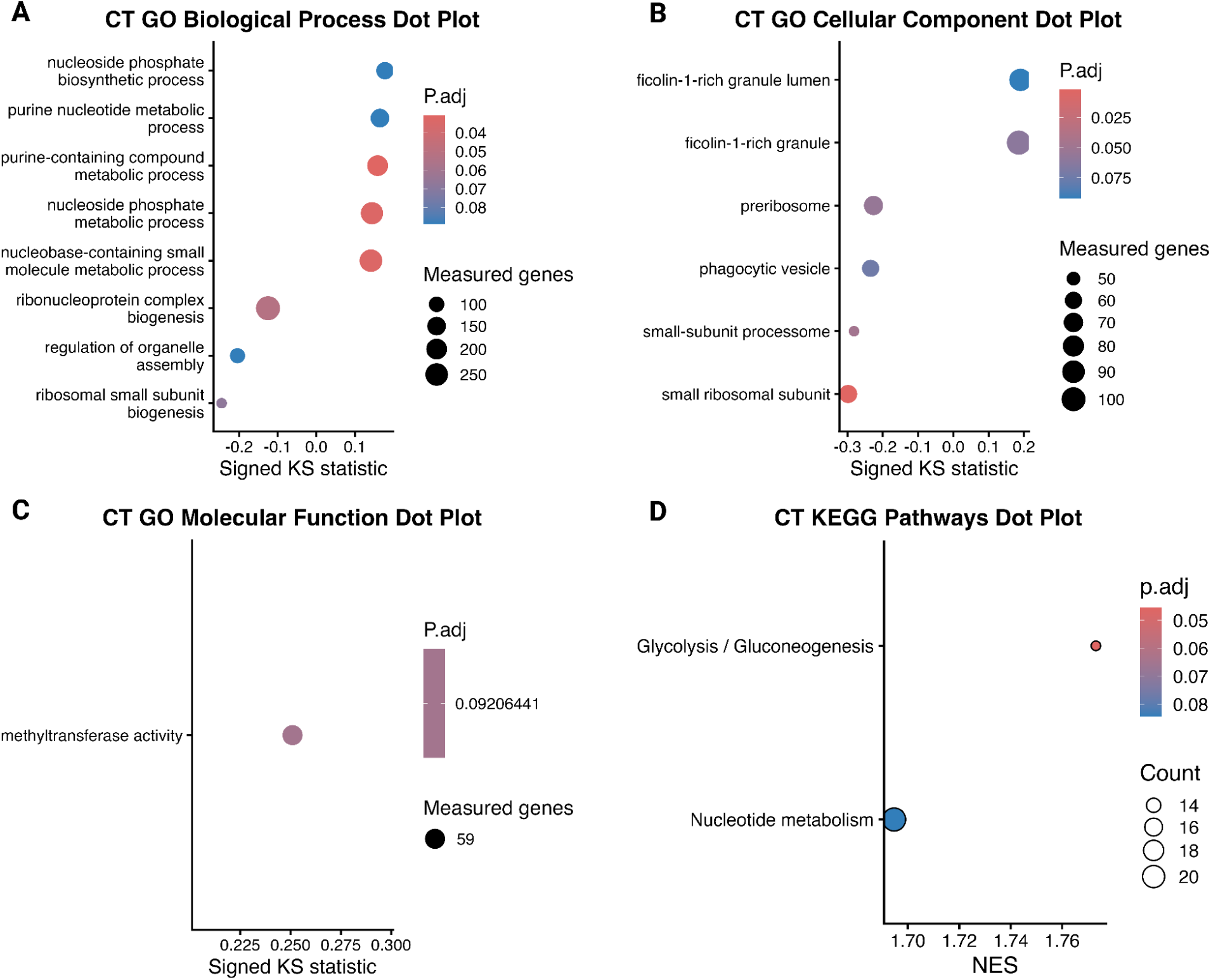
Pathway enrichment of conjunctive targeting. **A.** GO biological process analysis showing positive enrichment of nucleotide-related metabolism and negative enrichment of ribonucleoprotein complex and small-subunit biogenesis and organelle assembly. **B**, GO cellular component analysis showing positive enrichment of ficolin-1-rich granule compartments and negative enrichment of ribosome-associated compartments and the phagocytic vesicle. **C**, GO molecular function analysis identifying positive enrichment of methyltransferase activity. **D**, KEGG analysis showing positive enrichment of glycolysis/gluconeogenesis and nucleotide metabolism. For a–c, horizontal position indicates the signed KS statistic, and dot size represents the number of measured proteins assigned to each term. For d, horizontal position indicates the normalized enrichment score, and dot size represents the leading-edge protein count. Color indicates the adjusted P value. Conjunctive targeting (CT) terms were significant under combination treatment but nonsignificant or unavailable under both single-agent treatments.

KEGG enrichment provided additional evidence of metabolic involvement, with positive enrichment of glycolysis/gluconeogenesis (adjusted *P* = 0.046) and nucleotide metabolism (adjusted *P* = 0.084) in the conjunctive targeting set (**Fig. 5D**). No negatively enriched KEGG pathways were identified under the applied criterion. Together with the nucleotide-related GO findings, these results suggest potential involvement of nucleotide and glucose metabolism in combination-associated proteome remodeling, although the limited number of enriched pathways warrants cautious interpretation. The corresponding R, U, and RU dot plots show shared chromatin changes, stronger ribosomal signatures in U and RU, and additional metabolic enrichment in RU. Together, these analyses nominate nucleotide metabolism and glycolysis/gluconeogenesis, coupled to negative ribosome-biogenesis signatures and altered granule or vesicle compartments. Neither single-agent profile showed corresponding glycolysis/gluconeogenesis enrichment under the applied criteria, highlighting a combination-associated metabolic accessibility signature in NOMO-1 lysates. This finding nominates glycolytic proteins for mechanistic investigation, although the accessibility measurements do not directly establish glycolysis induction or inhibition of oxidative phosphorylation by either single-agent or combined treatment.

The gene–term networks organize conjunctive targeting into positive nucleotide metabolism and glycolytic signatures, a negative ribosome biogenesis signature, and distinct granule and vesicle compartments (**Supplementary Fig. 2**). Purine and nucleoside phosphate terms shared many annotated proteins, indicating a strongly overlapping nucleotide-associated signature. Ribonucleoprotein complex and small ribosomal subunit biogenesis formed a second connected group, consistent with their negative enrichment (**Fig. 5A**). In the cellular component network, the small ribosomal subunit, processome, and preribosome clustered separately from ficolin-1-rich granules and phagocytic vesicles (**Supplementary Fig. 2B**). The KEGG network separated glycolysis/gluconeogenesis from nucleotide metabolism; their 13 and 21 leading-edge proteins, respectively, did not overlap (**Supplementary Fig. 2D**). Across treatments, RU retained the chromatin response seen with each drug and the ribosomal features prominent under ulixertinib, alongside the conjunctive targeting-associated metabolic terms. Thus, the metabolic and ribosome-related findings involve distinct but internally overlapping sets of protein annotations.

### Peptide-level profiles reveal broad combination-associated accessibility changes

To determine whether combination-associated accessibility changes extended beyond the cutoff-defined target set, we compared peptide-level nominal *P*-value distributions across treatment conditions. Among 36,455 peptides quantified in all three conditions, the combination showed the greatest excess of small *P* values (**Fig. 6A**). Storey’s method^22^ estimated non-null peptide fractions of 22.1% for the combination, 21.4% for ulixertinib, and 12.4% for ruxolitinib. Using the prespecified criterion (nominal *P* < 0.05 and |log₂ fold change| > 1), 2,069 peptides showed altered accessibility under combination treatment, compared with 1,797 ± 30 under within-peptide treatment-label permutation, corresponding to a 1.15-fold excess (**Fig. 6B**). Ulixertinib and ruxolitinib yielded 1,943 and 1,380 peptides meeting the same criterion, respectively. Together, the full *P*-value distributions and threshold-based permutation analysis indicate that combination treatment is associated with widespread changes in peptide-level accessibility.

**Fig. 6.**
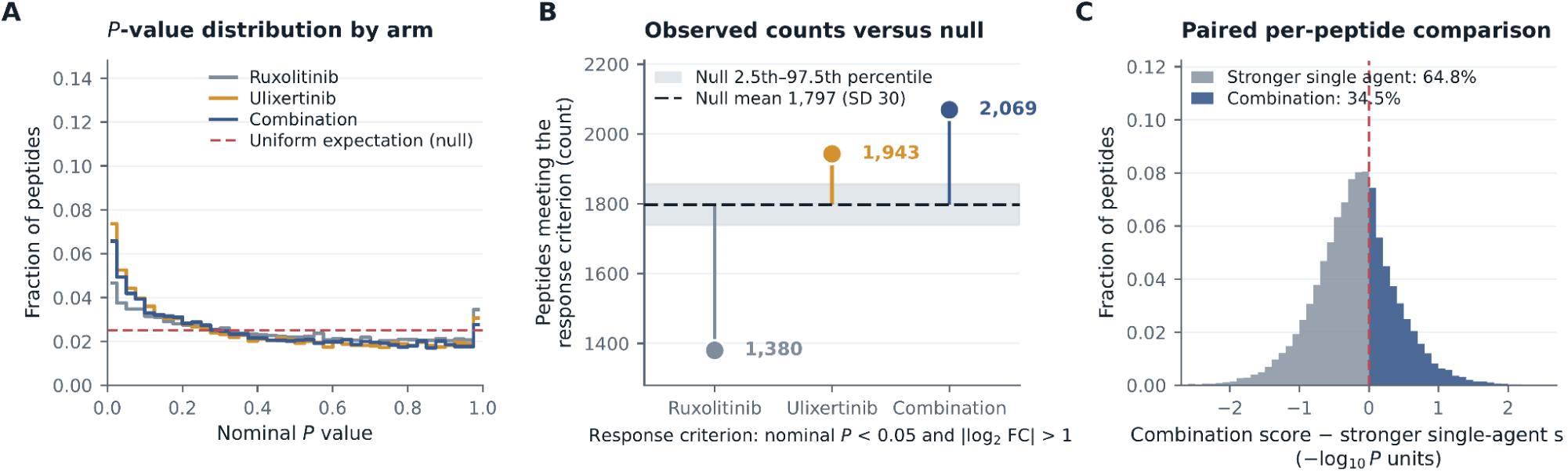
Global comparison of peptide accessibility changes across treatment conditions. **A.** Nominal P-value distributions across 36,455 peptides; dashed line, uniform expectation. **B**. Observed numbers of peptides with altered accessibility versus the exchangeability null. **C.** Peptide-wise comparison of combination-associated accessibility changes with those of the stronger single agent.

### Combination-associated accessibility changes are predominantly regional

Combination-associated changes were predominantly regional within proteins (**Fig. 7**). Among 211 conjunctive targeting candidates with at least four quantified peptides, 209 (99.1%) exhibited regional accessibility changes, whereas only two (0.9%) showed coordinated protein-wide changes (**Fig. 8**). The median fraction of peptides meeting the combination-associated accessibility-change criterion was 12%, indicating that altered accessibility typically involved a minority of the peptides quantified for each protein. Among the remaining peptides, the median across proteins of the absolute median signed log₂ fold change was 0.12. Thus, conjunctive targeting was generally characterized by altered accessibility in selected peptide regions against a background of comparatively small changes elsewhere in the protein, rather than by coordinated changes across its quantified peptides.

**Fig. 7.**
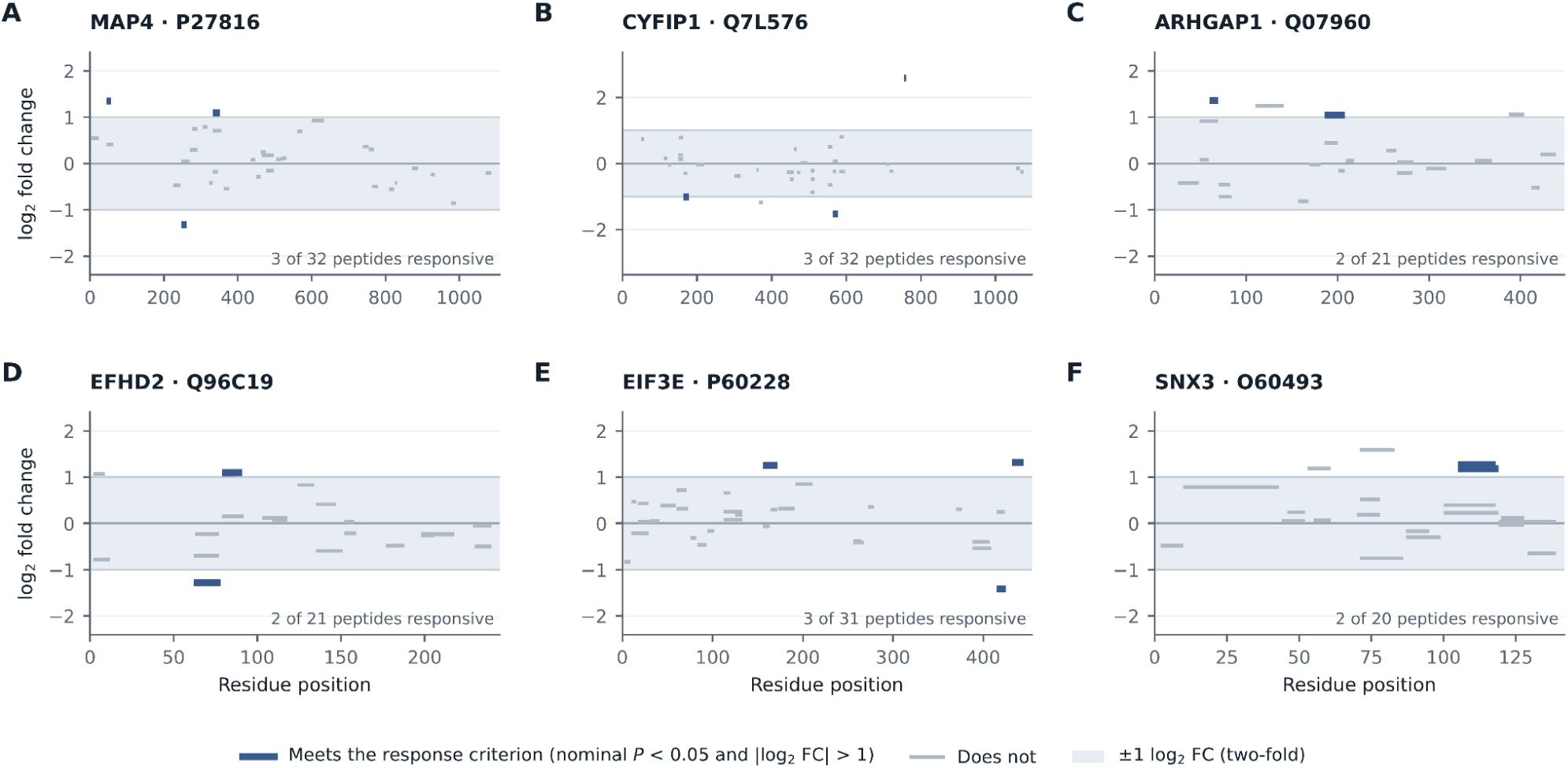
Peptide-level accessibility profiles of representative conjunctive targets. Indigo, peptides with altered accessibility; grey, peptides not meeting the prespecified accessibility-change criterion. Shading indicates the two-fold change boundary.

**Fig. 8.**
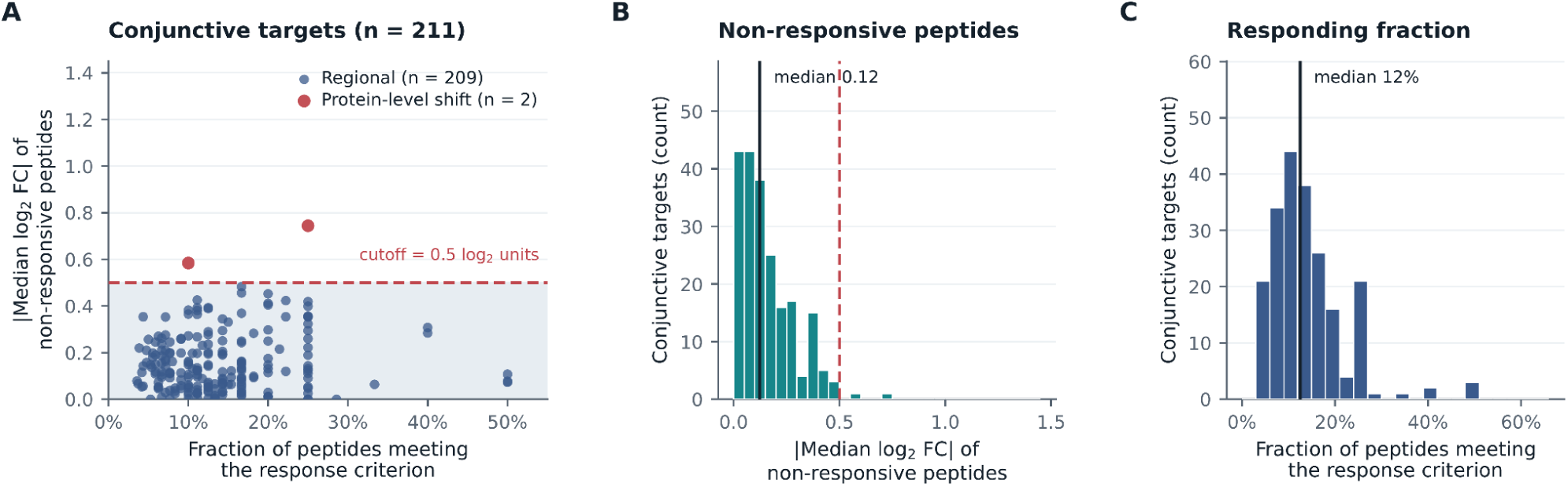
Combination-associated accessibility changes are predominantly regional. **A**, Fraction of peptides meeting the combination-associated accessibility criterion and absolute median log₂ fold change of the remaining peptides across 211 conjunctive targeting candidates. **B** and C are corresponding marginal distributions. Dashed lines indicate the predefined thresholds distinguishing regional accessibility changes from coordinated changes across peptides within the same protein.

Representative profiles further illustrated this pattern, with peptides showing altered accessibility occurring alongside peptides with comparatively small changes from the same protein (**Fig. 7**). Overall, these results support interpretation of combination-associated effects at peptide-level resolution and are consistent with regional changes in proteolytic accessibility rather than coordinated changes across peptides throughout the protein, a distinction that could be obscured by protein-level summarization.

### A Phospho (STY)-included search preserves the combination-response calls

The predominantly regional pattern of combination-associated peptide accessibility raised the hypothesis that local post-translational modifications (PTMs) may contribute to, or coincide with, these changes. Because the relevant PTM class was not known a priori, we applied the PTM-guided strategy described in the PEIMAN2 workflow to identify PTM annotations enriched among proteins exhibiting RU-associated accessibility changes^23^. This analysis highlighted phosphoprotein, phosphoserine, phosphothreonine, and phosphotyrosine annotations (**Fig. 9A**), pointing to phosphorylation as a candidate feature associated with the observed regional accessibility patterns. We therefore performed a second database search allowing phosphorylation on serine, threonine, and tyrosine [Phospho (STY)] to examine this possibility directly.

**Fig. 9.**
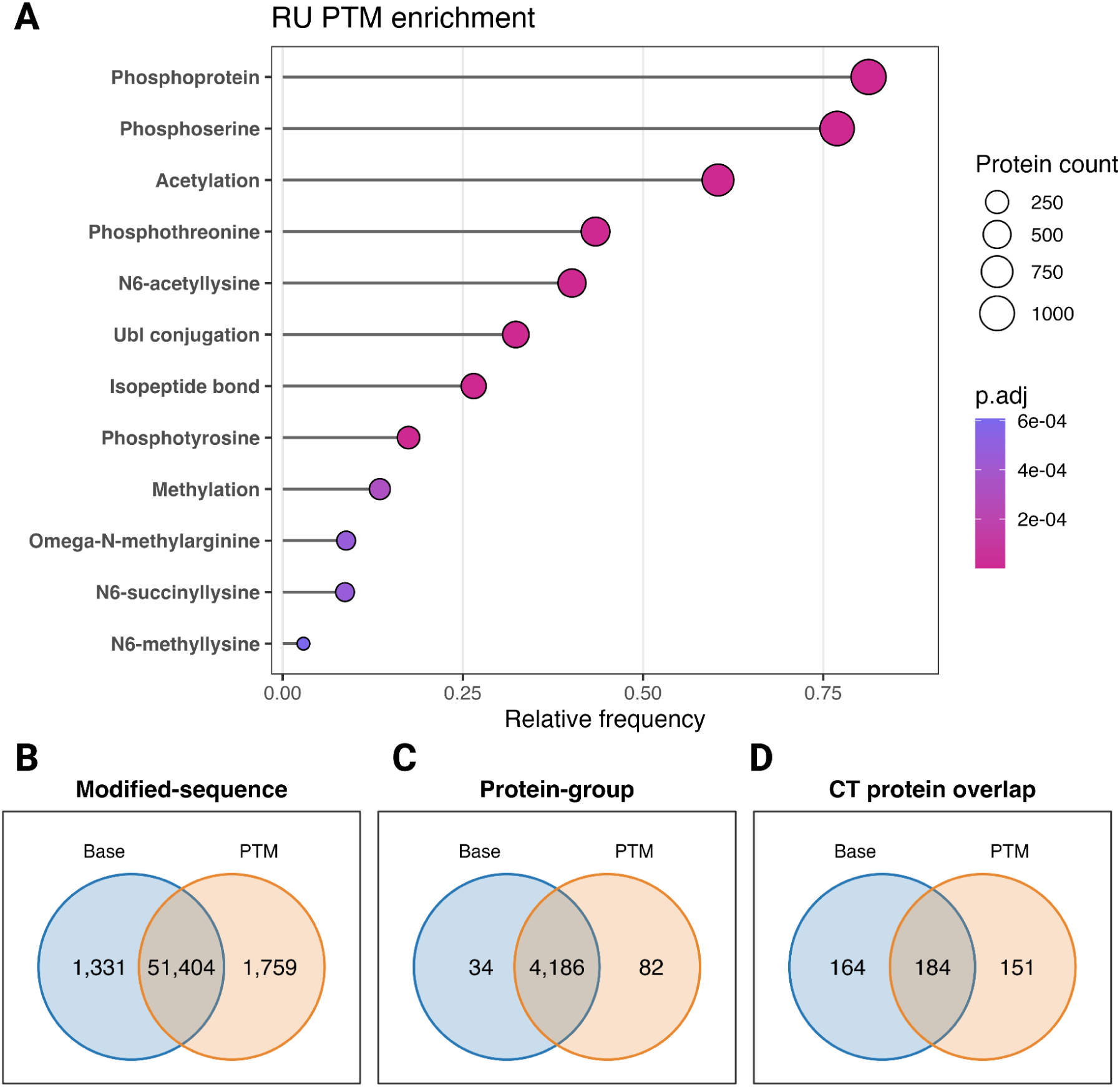
PTM-guided analysis and concordance of conjunctive targeting calls. **A.** PEIMAN2 analysis of PTM annotations enriched among proteins with RU-associated accessibility changes. Position indicates relative annotation frequency, point size indicates the number of proteins, and color indicates the adjusted P value. Concordance between the Base and Phospho (STY)-included searches at the level of exact modified-peptide sequences (**B**) and protein groups (**C**) under matched identification-quality criteria. **D.** Concordance of conjunctive targeting (CT) protein calls between the corresponding downstream analyses. Base denotes the original search, and PTM the Phospho (STY)-included search. CT denotes proteins meeting the predefined criteria for combination-associated accessibility changes but not the corresponding single-agent criteria.

Under matched identification-quality filters, the Phospho (STY)-included search yielded 8,364 Phospho (STY)-annotated MS2 evidence rows. These represented 247 peptide sequences, 384 modified-sequence forms, and 164 exact protein groups; 78 of the sequences were absent from the base search. Across the searches, 42,005 peptide sequences and 4,186 exact protein groups were shared, retaining 98.0% and 99.2% of the respective base-search sets (protein-group overlap, **Fig. 9C**). The modified-sequence comparison identified 51,404 shared forms (**Fig. 9B**). As expected, the expanded search recovered additional modification-annotated evidence while largely preserving the overall identifications.

The downstream CT analysis identified 348 proteins in the base search and 335 in the Phospho (STY)-included search, with 184 shared calls (**Fig. 9D**). PTPN11, encoding SHP2, was among the shared proteins, although its retained evidence was not Phospho (STY)-annotated. SHP2 is relevant to the ERK-directed component of RU: in FLT3-ITD AML models, feedback through SHP2 restored ERK signaling during SHP2 inhibition.^24^ EIF4EBP1 entered the CT set only in the Phospho (STY)-included search and had a Phospho (STY)-annotated sequence absent from the base search. Studies in AML have linked 4E-BP1 phosphorylation to translational control and shown an interaction between ERK activity, 4E-BP1 phosphorylation and cell survival^25,26^. These two proteins therefore offer focused candidates for testing how RU affects signaling and translation.

## Discussion

CoPPRA extends the analysis of drug combinations from proteome-wide changes to regional differences in protein accessibility. Established structural proteomics approaches, including LiP-MS and PELSA, use limited proteolysis to capture treatment-associated changes in protein structure and proteolytic accessibility at peptide-level resolution ^7,15^. Building on these principles and the high-ratio partial proteolysis strategy used in HOLSER, CoPPRA adapts this framework to the comparative analysis of single and combined drug treatments^16^. This design enables combination-associated effects to be localized to specific peptide regions, providing structural context complementary to protein-level stability and solubility measurements obtained with approaches such as thermal proteome profiling (TPP)^8^, Proteome Integral Solubility Alteration (PISA)^8,27^, and its combinatorial extension, CoPISA^11^.

CoPPRA adds regional resolution to the conjunctive targeting signature previously detected by CoPISA through changes in protein solubility and stability^11^. In NOMO-1 cell lysates, ruxolitinib plus ulixertinib produced peptide accessibility changes only under combined treatment. These regions give the combination response a concrete structural context and focus mechanistic investigation on how the two compounds jointly influence local protein states (AND-gate logic). The substantial ulixertinib-associated response and heterogeneous peptide-level effects under RU make the identity and location of responsive regions particularly informative for understanding the combination. Mapping these regions onto protein domains and interaction interfaces can guide tests of whether combined exposure alters local conformation, ligand binding, or complex assembly. Proteolytic accessibility can report both direct engagement and indirect changes in protein state,^7,9^. so the implicated regions provide experimentally tractable starting points for resolving the physical basis of conjunctional targeting.

The distribution of accessibility changes within individual proteins further supports a predominantly regional interpretation. In most conjunctive targeting candidates, altered accessibility was confined to a subset of quantified peptides, while the remaining peptides showed comparatively small changes. Conjunctive targeting therefore describes a peptide-level accessibility pattern rather than a demonstrated change across the entire protein or in total protein abundance. These regional profiles provide candidate protein regions for subsequent structural and functional validation.

The enrichment of nucleotide metabolism, glycolysis/gluconeogenesis and ribosome biogenesis places the combination-associated accessibility response at the intersection of metabolic supply and biosynthetic demand. This convergence is relevant to AML, where nucleotide availability and ribosome biogenesis support proliferative and metabolic adaptation^28–30,31^. The metabolic and ribosomal convergence is mechanistically plausible, while the granule and vesicle terms offer a compartmental context. Enrichment direction reflects accessibility-associated rankings, not measured pathway flux or proof that RU activates nucleotide synthesis or suppresses ribosome production.

The combination-associated glycolysis/gluconeogenesis signature raises the possibility that joint JAK–ERK inhibition affects metabolic protein states beyond the responses detected with either drug alone. This is relevant to AML, where oxidative metabolism supports chemotherapy-resistant cells, and guanine nucleotide availability can sustain ribosome biogenesis and glycolytic reprogramming^3,30^. With the shared mitochondrial candidates OPA1 and STOML2, the enrichment motivates testing whether RU alters mitochondrial organization and the balance between glycolytic and mitochondrial energy metabolism. Positive accessibility enrichment, however, does not specify metabolic flux, and the present data do not demonstrate OXPHOS inhibition or compensatory glycolysis. Measuring oxygen consumption, glycolytic proton efflux, and isotope-resolved glucose metabolism after single-agent and combination treatment in intact AML cells would establish whether these structural signatures correspond to metabolic adaptation or a vulnerability contributing to drug sensitivity.

22 proteins were shared between the CoPPRA and CoPISA conjunctive targeting sets in NOMO-1 cell lysates, linking candidates detected through complementary measurements of peptide accessibility and protein solubility.^11^ OPA1 provides a direct link to AML biology, as its regulation of cristae structure contributes to venetoclax resistance^32^. STOML2’s role in respiratory-chain supercomplex formation further motivates examining whether mitochondrial organization connects the structural response to altered respiratory function.^33^ The ribosomal, RNA-export, genome-maintenance and proteostasis candidates broaden this interpretation to the production and maintenance of cellular macromolecules. Their biological significance lies in the possibility that perturbing several of these functions together could reduce the capacity of leukemic cells to tolerate treatment. This provides a testable model in which conjunctive targeting creates vulnerabilities through changes in protein states across multiple cellular processes. Agreement between complementary biophysical measurements helps prioritize the proteins through which to investigate this model, while CoPPRA’s regional resolution directs functional experiments toward the responsive peptide regions.

Ruxolitinib inhibits JAK1/2 signaling, whereas ulixertinib targets ERK1/2 within the MAPK pathway^17,18^. The shared candidates therefore motivate testing whether combined JAK–ERK inhibition affects mitochondrial organization, RNA processing, genome maintenance, and protein quality control in AML cells. Their detection by both assays strengthens their priority for mechanistic investigation, although the lysate measurements do not establish downstream signaling effects or direct drug binding. By adding regional accessibility information to CoPISA-derived candidates, CoPPRA provides a basis for selecting protein regions and functional assays to test how combination-associated structural changes contribute to drug sensitivity.

The PTM analysis adds a complementary view of the RU response by connecting responsive proteins to prior phosphorylation annotations and bringing modified peptides into candidate discovery. Differences between the base and expanded searches show that candidate selection depends partly on the evidence included in the search. The results therefore help prioritize signaling and translational-control proteins for validation.

Establishing treatment-regulated phosphorylation or a phosphorylation-dependent conjunctive targeting mechanism will require localized phosphosites and direct measurements of their treatment response.

The study also has clear limitations. CoPPRA was performed in non-denaturing lysates, which preserves many protein-state features but cannot reproduce intact-cell compartmentalization, active metabolism, drug transport, or microenvironmental interactions. Partial proteolysis reports regional accessibility indirectly and does not by itself establish direct binding, causal pathway regulation, or atomic structural change^7,9^. The design also contains three technical and three biological replicates per condition, and threshold-defined CT membership remains sensitive to statistical cutoffs, as illustrated by proteins moving across the significance boundary between searches. Although the combination produced a broad peptide response, its absolute change was often smaller than that of the stronger single agent. Ulixertinib produced a larger single-agent response than ruxolitinib, while the contribution of each drug to the combination remains unresolved. Treating intact AML cells before rapid lysis, alongside matched protein abundance measurements, could test which treatment-associated accessibility patterns remain detectable after extraction.

## Methods

### Cell culture

NOMO-1 cell lines from Deutsche Sammlung von Mikroorganismen und Zellkulturen (DSMZ, Germany) were cultured in RPMI-1640 medium containing L-glutamine (Gibco, Thermo Fisher Scientific, USA) plus 10% fetal bovine serum (FBS) (Gibco, Thermo Fisher Scientific) and 1% penicillin/streptomycin (Gibco, Thermo Fisher Scientific) at 37°C and 5% CO₂.. Cells were harvested and centrifuged at 600 g for 5 minutes.

### CoPPRA pipeline

Cells were harvested at 1.5-2M/mL to ensure they are in optimal density for growth. All cell lines were tested for mycoplasma contamination using PCR-based assays to confirm the absence of infection. For native protein extraction, NOMO-1 cells were collected and washed twice with ice-cold PBS, followed by resuspension in 20 mM EPPS (pH 8.2) containing EDTA-free protease inhibitors. Cells were disrupted by three freeze-thaw cycles, and insoluble material was removed by centrifugation (16,000 × g, 10 min, 4°C). The resulting supernatants were used as native lysates for subsequent CoPPRA analysis. Protein concentration was quantified by bicinchoninic acid (BCA) assay, and samples were adjusted to 1 mg/mL before downstream treatment.

For CoPPRA analysis, native protein lysates were prepared from NOMO-1 cells in 20 mM EPPS buffer (pH 8.2) supplemented with 1x EDTA-free protease inhibitors (Roche, Switzerland), followed by clarification by centrifugation. Protein concentrations were determined by BCA assay, and all lysates were normalized to 1 mg/mL before downstream processing. For each treatment condition, 50 µg of NOMO-1 native lysate was incubated for 30 min at 25°C with ruxolitinib, ulixertinib, the combination, or vehicle control under matched solvent conditions (final drug concentration 3 µM, DMSO concentration 0.06%). Each treatment condition was prepared in three biological replicates. Additional fully digested reference samples were prepared and analyzed in separate LC-MS/MS runs. After drug incubation, treatment lysates were subjected to short native trypsin pulse proteolysis at a 1:1 enzyme-to-protein ratio for 4 min at 25°C to capture ligand- and combination-dependent differences in regional protein accessibility^7,9^. In parallel, these reference samples were digested for 8 h to generate peptide material for identification.

Proteolysis was quenched by adding 8 M guanidine hydrochloride prepared in 20 mM EPPS to a final concentration of 6 M. Proteins and peptides were reduced with 10 mM TCEP and alkylated with 40 mM iodoacetamide for 45 min at room temperature in the dark. The samples were subsequently diluted with EPPS buffer to reduce the guanidine hydrochloride concentration to approximately 1 M, acidified to pH 2-3 with trifluoroacetic acid, and desalted individually using C18 cartridges. The purified peptide samples were dried by vacuum centrifugation and stored until LC-MS/MS analysis.

### LC-MS/MS analysis

Prior to LC–MS/MS analysis, 200 µL of 0.1% formic acid (FA) was added to each sample tube. Subsequently, 15 µL of each peptide digest solution was loaded onto Evotips (Evosep ApS) according to the manufacturer’s instructions. Peptide samples were analyzed using an Evosep Eno LC system (Evosep ApS) operated with the 60 samples per day (60 SPD) gradient method. Chromatographic separation was performed using an 8 cm × 150 µm, 1.5 µm particle size. Performance-analytical column (Evosep, #EV1109) maintained at 40°C. Mobile phase A consisted of 0.1% formic acid in water, while mobile phase B consisted of 0.1% formic acid in acetonitrile. The LC system was coupled online to an Orbitrap Astral mass spectrometer (Thermo Fisher Scientific) equipped with an EASY-Spray source and controlled using Thermo Scientific Xcalibur software (version 4.7.102.25). Each sample was analyzed in three technical replicate runs using the same LC–MS/MS acquisition method.

Mass spectrometric analysis was performed in positive-ion data-independent acquisition (DIA) mode. MS1 spectra were acquired at a resolution of 240,000 over an m/z range of 380–980. The RF lens was set to 40%, the MS AGC target was set to 500%, and the maximum injection time was 5 ms. DIA MS/MS spectra were acquired using a 2 Th isolation window over an m/z range of 150–2000. The MS/MS AGC target was normalized to 500%, and higher-energy collisional dissociation (HCD) was performed using a normalized collision energy of 25%.

### Peptide spectra matching and quantification

Raw LC-MS/MS output files were processed using MaxQuant version 2.8.1.0 with the MaxDIA workflow^34,35^. Spectra were searched against the human UniProt reference proteome sequence database (UP000005640; taxonomy ID 9606, downloaded 16.07.2026)^36^, with 26,785 protein entries, supplemented with reversed sequences for false discovery rate estimation^37^. Searches were performed with Trypsin/P specificity, allowing a maximum of 5 missed cleavages. Carbamidomethylation of cysteine was set as a fixed modification, while methionine oxidation and protein N-terminal acetylation were included as variable modifications. The minimum peptide length was seven amino acids, the match between runs was included, and peptide- and protein-level false discovery rates were controlled at 1% or less.

Intensity values were extracted from the MaxQuant evidence-level output. Reverse identifications, potential contaminants, entries without positive intensity values, and evidence rows assigned to more than one MaxQuant protein group were excluded. Within each raw file, repeated evidence entries corresponding to the same modified peptide sequence and precursor charge were summarized by the median log₂ intensity.

Normalization was performed at PSM level. Within each biological experiment, precursors detected across all three technical runs were used to calculate the raw-file medians, and each run was median-centered accordingly. The normalized precursor intensities were then summarized to peptide level by median aggregation. Technical runs were subsequently combined at the biological experiment level by taking the median normalized intensity for each peptide. Peptides were retained when detected in at least 20% of raw files. Missing values in the matrix were replaced with the median observed abundance of the corresponding peptide.

Reference samples were normalized and evaluated separately but excluded from treatment reproducibility metrics and differential accessibility testing. For the primary biological analysis, peptides were required to be quantified in at least two of the three biological replicates in each treatment condition. Differential peptide accessibility was assessed using the limma package (version 3.68.4)^38^, with treatment contrasts defined relative to the vehicle control. Differentially accessible peptides were subsequently mapped to MaxQuant protein groups, expanded to their constituent UniProt accessions, and consolidated into nonredundant protein-level summaries for downstream functional analysis. To determine whether the observed number of peptides with altered accessibility exceeded that expected by chance, condition labels were randomly permuted across the 12 biological samples 10,000 times while preserving three samples per condition. Each permutation was applied jointly across all peptides to preserve the correlation structure among peptide measurements^39–41^.

### Peptide-resolved profiling and sequence mapping of selected proteins

Peptide-level statistics and normalized biological-replicate intensities for MEK1/MAP2K1 and ATP6V1G1 were extracted from the limma CoPPRA analysis. All retained peptide sequences assigned to these proteins were included, irrespective of statistical significance. The ATP6V1G1 protein sequence and positional annotations were retrieved from UniProtKB (O75348; accessed 24.09.2026)^36^. Unmodified peptide sequences were mapped by exact sequence matching, with positions reported as one-based, inclusive coordinates relative to the full-length sequence. Mapping ambiguity was assessed against 42,562 reviewed human UniProtKB canonical and isoform sequences downloaded on the same date. Exact matches were recorded, and additional searches treating isoleucine and leucine as equivalent were used to identify potential ambiguity. Matches to other genes and alternative isoforms were recorded separately. UniProt-annotated α-helices were displayed alongside the mapped peptide intervals. MEK1 peptide responses were visualized as a heatmap, and selected MEK1 and ATP6V1G1 peptides were displayed using individual biological-replicate intensities and group medians. ATP6V1G1 sequence maps included all retained peptides under each treatment condition. Heatmaps, replicate plots, and sequence maps used a common color scale spanning log₂FC values from −5 to 5.

### Functional enrichment and network analysis

Functional enrichment and network analyses were performed in R using topGO (version 2.58.0)^42^, clusterProfiler (version 4.14.6)^43^ and enrichplot (version 1.26.6). Peptide-level limma statistics were converted to signed normal scores according to nominal *P* values and log₂ fold-change direction, summarized by UniProt accession, and mapped to Entrez gene identifiers and aggregated by median to generate ranked gene lists for ruxolitinib, ulixertinib and combined treatment.

GO biological process, cellular component, and molecular function enrichment^44,45^ was assessed using the classic topGO Kolmogorov–Smirnov test in both rank directions. Directional *P* values were combined as min(1, 2 × min(*P*up, *P*down)) and adjusted using the Benjamini–Hochberg method^46^. Human KEGG pathways^47,48^ were analyzed by GSEA using fgsea^49,50^, with normalized enrichment score (NES) describing enrichment direction. Gene sets containing 15–500 measured genes were tested, and GO terms were tested for enrichment toward both ends of the ranked gene list. The resulting *P* values were adjusted by Benjamini–Hochberg correction. Terms with adjusted *P* < 0.10 were retained. Conjunctive targeting terms were defined as significant in the combined treatment but nonsignificant or unavailable in both monotherapies^11^.

Dot plots displayed signed KS statistics for GO and NES for KEGG. Gene–term networks showed measured term members for GO and leading-edge genes for KEGG. Enrichment maps^51^ were based on Jaccard term similarity with a minimum similarity of 0.20, displaying up to 12 categories.

### Structural analysis of conjunctive targets

We used limma coefficients as log₂ fold changes so that effect sizes corresponded to the reported P values. In each treatment condition, a peptide was considered responsive at nominal P < 0.05 and |log₂ fold change| > 1. Among peptides quantified in all three treatment conditions, we compared each treatment condition with the stronger of the other two using −log₁₀(P), splitting ties fractionally. We repeated the comparison using |log₂ fold change| and estimated the unchanged fraction in each treatment condition by Storey’s method^22^. For an exchangeability null, we permuted treatment labels within each peptide 2,000 times and compared the observed response counts with the resulting distribution.

### Regional versus protein-level changes

For conjunctive targets with at least four quantified peptides and at least one combination-responsive peptide, we calculated the absolute value of the median *signed* log₂ fold change among non-responsive peptides. Values >0.5 indicated a protein-level shift; other proteins were classified as regional. Taking the median before the absolute value distinguishes a consistent shift from peptides moving in opposite directions.

### Pockets and peptide positions

We retained AlphaFold residues with pLDDT ≥70^52^. Pockets were retained when fpocket and P2Rank detected them within 8 Å of each other and the fpocket druggability score was ≥0.20^53–55^. Pocket pairs were considered separable when they were at least 12 Å apart, no farther apart than twice the radius of gyration, on the same chain, and had interpocket predicted aligned error < 5 Å. Peptides were mapped by exact sequence match, and distances were measured from peptide centroids to the nearest pocket center. For continuous descriptors, we ranked peptides within each protein and tested the mean normalized rank of responsive peptides against 0.5 by the Wilcoxon signed-rank test across proteins. Each protein contributed equally; intervals were bootstrap percentiles over proteins. Spatial clustering was tested against 2,000 within-protein permutations of response labels. Pocket-detection thresholds were fixed before cohort analysis.

### Modification sites and interfaces

Curated modifications, binding, and active sites came from UniProt^36^. In experimental structures, interfaces comprised residues within 5 Å of another polymer chain of at least 20 residues, provided at least 10 residues participated in the contact. Structure positions were mapped to UniProt by sequence alignment requiring ≥80% identity. Because site annotations were sparse, peptide site overlap was analyzed with protein-stratified Mantel–Haenszel odds ratios and Robins–Breslow–Greenland intervals^56,57^.

### Post-translational modification enrichment analysis

Post-translational modification (PTM) enrichment analysis was performed using human annotations from PEIMAN2 (version 1.0.2)^23^. UniProt accessions were matched to the bundled database. Significant proteins were defined by at least one peptide meeting the differential accessibility criteria for each treatment. SEA assessed PTM over-representation in the R, U and RU protein sets using a one-sided hypergeometric test against the mapped protein background. PSEA ranked the complete mapped background by median peptide limma log₂ fold change and used the fgsea multilevel algorithm (version 1.32.4)^49^ to assess directional enrichment. For both analyses, categories without significant annotated protein hits were excluded before Benjamini–Hochberg correction^46^ within each treatment comparison. Terms with adjusted *P* < 0.05 were retained, with fold enrichment > 1 additionally required for SEA. PSEA retained positive and negative enrichment, while reported leading-edge proteins were restricted to those significant in the corresponding treatment. The ranked protein background remained unchanged by this reporting filter.

To evaluate modification-associated identifications and the robustness of the conjunctive targeting results, the raw data were searched a second time using a Phospho (STY)-included MaxQuant configuration. The same overall search and downstream processing workflow was retained, with Phospho (STY) included as an additional variable modification. The resulting evidence was processed using the same filtering, normalization, peptide aggregation and limma differential-accessibility workflow as the base search. Concordance between the base and Phospho (STY)-included searches was evaluated at the levels of unique peptide sequence, modified-sequence form, and exact MaxQuant protein group.

Conjunctive targeting proteins were defined independently in each search as proteins showing significant accessibility changes under combined ruxolitinib–ulixertinib treatment but not under either single-agent treatment. Protein accessions were matched exactly between searches, and the intersection of the two CT sets was defined as the common CT set for subsequent functional analysis. Modified evidence from the Phospho (STY)-included search was additionally mapped back to CT proteins to identify modification-bearing candidates and to determine whether the corresponding modified peptide sequences themselves showed significant RU-associated accessibility changes.

## Data Availability

The mass spectrometry proteomics data generated in this study have been deposited in the ProteomeXchange Consortium via the PRIDE partner repository under accession code PXD084605 [http://proteomecentral.proteomexchange.org/cgi/GetDataset?ID=PXD084605]. Source data are provided with this paper. There are no restrictions on data availability.

## Code Availability

The analysis code used in this study is publicly available at github.com/jafarilab/CoPPRA_Code.

## Acknowledgements

This study was financially supported by the Tampere Institute for Advanced Study and the Jane and Aatos Erkko Foundation [Grant 220031 to M.J.] and the Faculty of Medicine, University of Helsinki, Finland. A.A.S. acknowledges funding from the Swedish Research Council (2023-02692) and the Swedish Cancer Society (24 3595 Pj). Proteomics analyses were conducted at the Meilahti Proteomics Unit (supported by HiLIFE and Biocenter Finland).

## Author Contributions Statement

M.J. conceived the study and formulated the research questions. M.J. and A.A.S. provided scientific direction, supervised the project, and contributed to manuscript preparation. G.C. led the experimental analyses and interpretation of the experimental data, with E.G. contributing to experimental analyses and sample preparation. R.S. performed the proteomics mass spectrometry experiments and associated data analysis. P.B. designed the structural and statistical analyses and, together with M.M., developed the computational workflows, conducted the analyses, interpreted the results, and contributed to drafting and revising the manuscript. R.R. and M.V. contributed to the interpretation of the findings and provided critical input on the manuscript.

## Supplementary Figures

**Supplementary Fig. 1.**
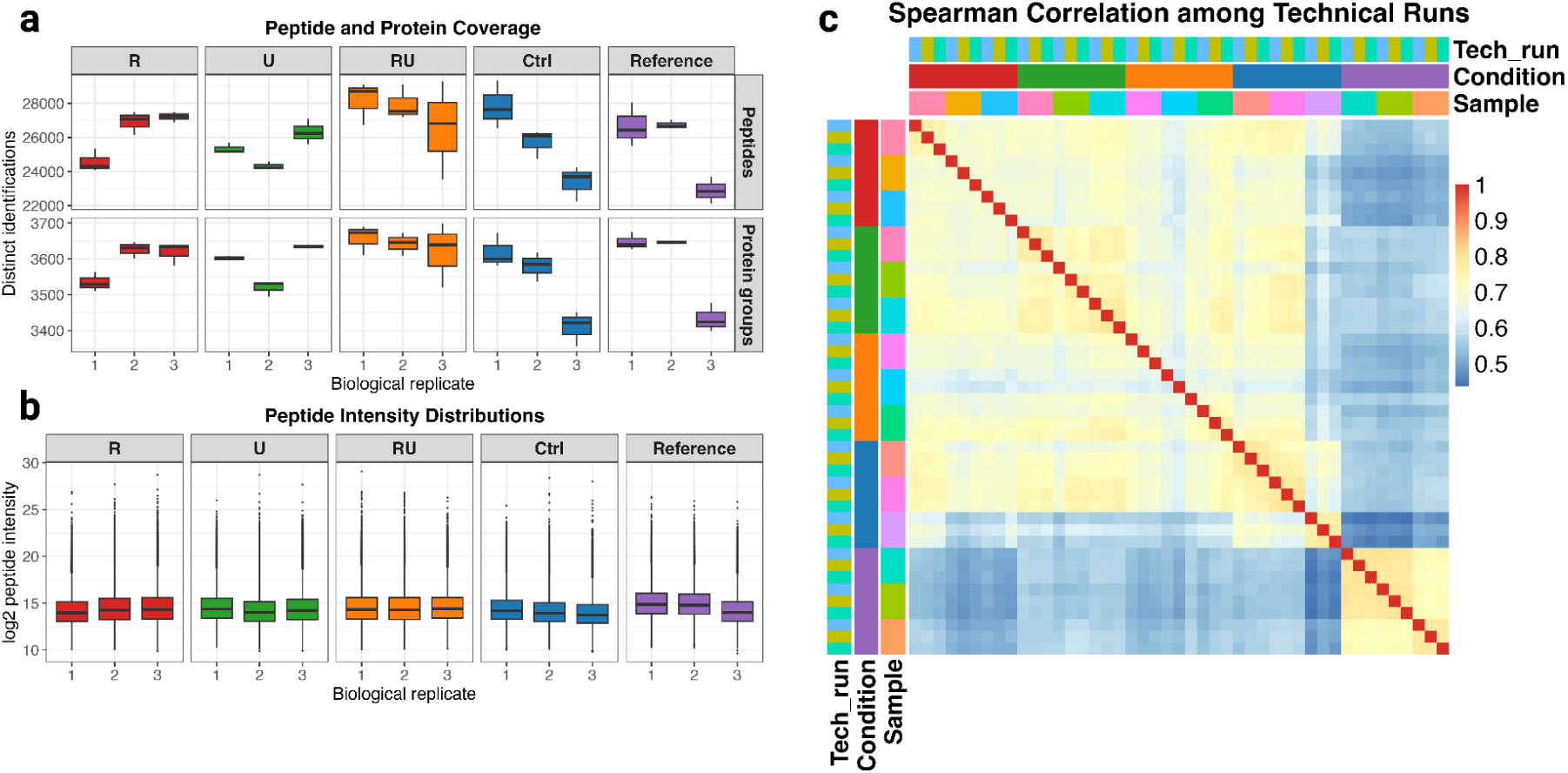
Quality-control assessment of peptide-level CoPPRA measurements. **A.** Distributions of unique peptide sequence counts (top) and leading protein group counts (bottom) across technical runs for three biological replicates per treatment condition and the fully digested reference samples (Reference in the plots). **B.** Distributions of log₂-transformed peptide intensities for each biological replicate, grouped by treatment condition or fully digested reference sample. R, ruxolitinib; U, ulixertinib; RU, ruxolitinib plus ulixertinib; Ctrl, vehicle control. **C**. Pairwise Pearson correlation heatmap of peptide intensities across technical runs, with annotations indicating technical replicate, treatment condition, and biological sample.

**Supplementary Fig. 2.**
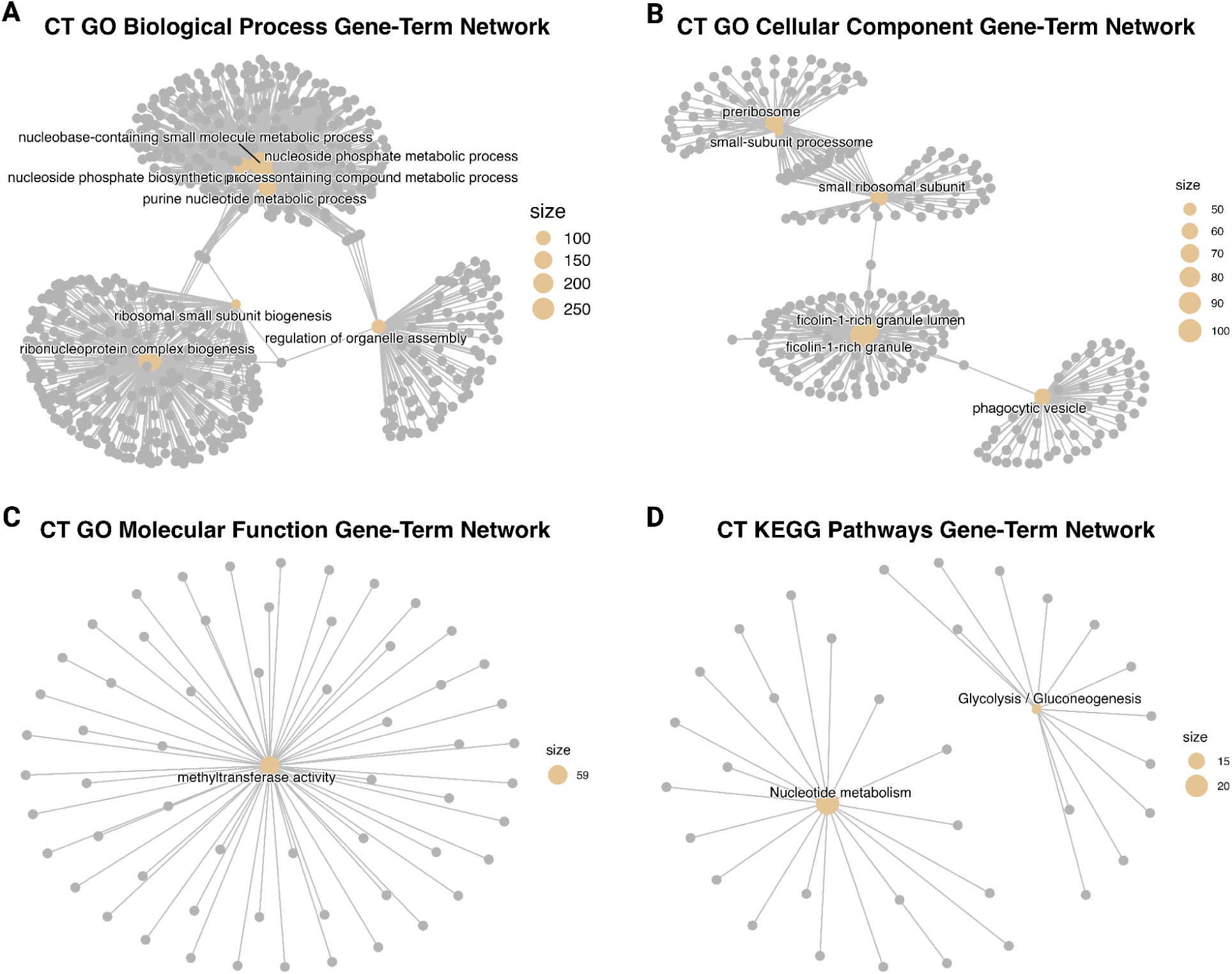
Network organization of conjunctive targeting pathways. **A**, GO biological process network showing an overlapping nucleotide-metabolism module, a ribonucleoprotein and small-subunit-biogenesis module, and an organelle-assembly branch. **B**, GO cellular component network showing ribosome-associated, ficolin-1-rich granule and phagocytic-vesicle groups. **C**, GO molecular function network centered on methyltransferase activity. **D**, KEGG network showing separate glycolysis/gluconeogenesis and nucleotide-metabolism modules with non-overlapping leading-edge gene sets. Gold nodes represent terms, whereas grey nodes represent mapped genes. The GO networks include all measured term members, whereas the KEGG network includes leading-edge genes. Gold-node size indicates the corresponding gene count, and edges indicate term–gene membership.

