## Supplementary figures and images for "Conjunctive Targeting Links Drug Synergy to Emergent Proteome Structural States"

### Supplementary figure 1

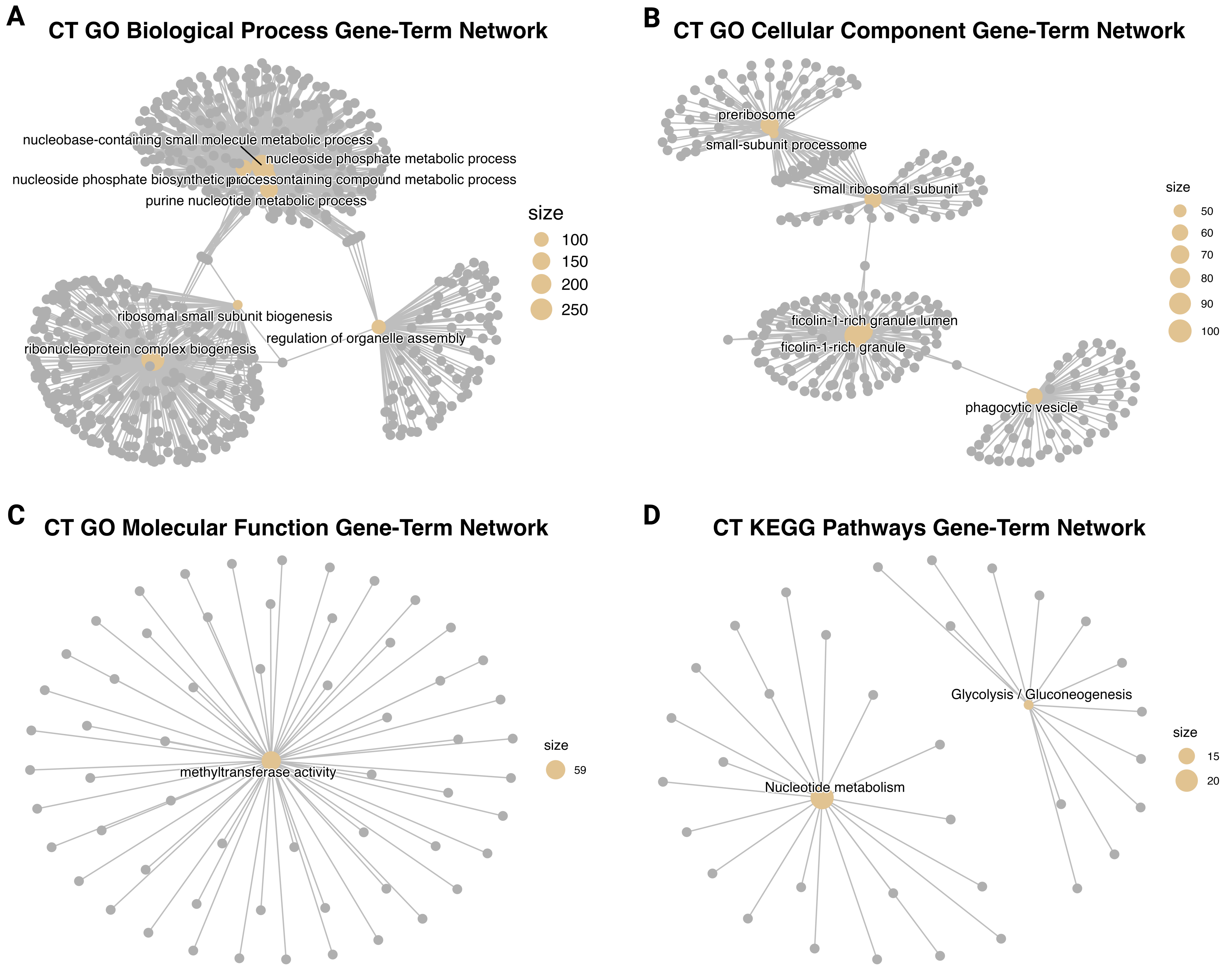

### Supplementary figure 2

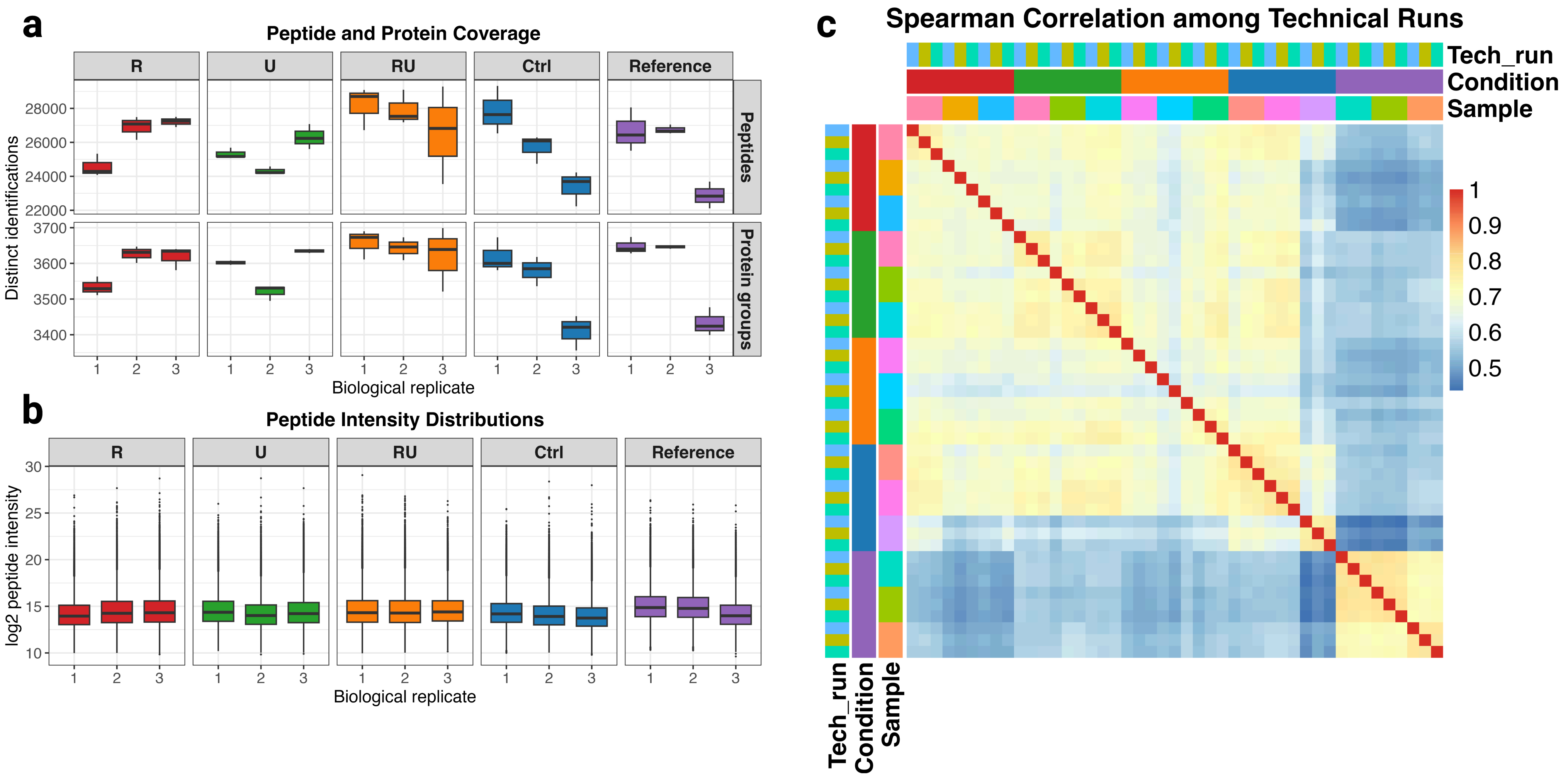
